# Lysyl oxidase-dependent bone marrow stiffening promotes neutrophil activation in diabetes

**DOI:** 10.1101/2025.08.30.673184

**Authors:** Mahesh Agarwal, Sathishkumar Chandrakumar, Vihar Trada, Samanvaya Srivastava, Kaustabh Ghosh

## Abstract

Persistent neutrophil activation contributes to the development of diabetic vascular complications in many organs including the retina. Past studies have implicated abnormal molecular or biochemical cues in neutrophil activation. Here we show that neutrophils can also be mechanically activated through an increase in bone marrow (BM) stiffness that results from diabetes-induced overexpression of matrix collagens and their crosslinker lysyl oxidase (LOX). Administration of LOX inhibitor β-aminopropionitrile (BAPN) in diabetic mice concomitantly blocked BM matrix stiffening and neutrophil activation (superoxide generation and gp91^phox^ upregulation), which predictably inhibited neutrophil cytotoxicity towards co-cultured retinal endothelial cells. Further, culturing neutrophils within BM-mimicking hydrogels of tunable stiffness indicate that matrix-localized and soluble LOX act synergistically to mechanochemically activate neutrophils in the BM. Thus, LOX-dependent changes in the BM physical environment offer a new mechanistic basis and therapeutic target for diabetic complications marked by neutrophil activation.

## Introduction

Diabetes leads to vascular complications in the retina, heart, kidneys, and brain. Neutrophil activation has been implicated in the pathogenesis of diabetes-induced vascular dysfunction (1, 2). But how neutrophils become activated in diabetes remains insufficiently understood. Neutrophils reside within the physical microenvironment of the bone marrow (BM) (3). A key component of the BM physical microenvironment is collagen-rich extracellular matrix that is amenable to crosslinking and stiffening by lysyl oxidase (LOX) (4, 5). We recently showed that an increase in LOX-mediated crosslinking of subendothelial collagen matrix increases retinal vascular stiffness in diabetes that, in turn, activates retinal endothelial cells (EC) and promotes their neutrophil-induced apoptosis, hallmarks of early diabetic retinopathy, a major vision-threatening retinal microvascular complication of diabetes (6, 7). Here we asked whether diabetes simultaneously leads to an increase in LOX-mediated BM matrix stiffness and, if so, whether the stiffer BM matrix mechanically activates neutrophils and renders them cytotoxic towards retinal ECs.

## Results and Discussion

### LOX promotes BM matrix stiffening and neutrophil activation in diabetes

Diabetes induces LOX upregulation in the retina and aorta where it enhances subendothelial matrix deposition and crosslinking (6, 8). To determine whether diabetes exerts similar effects on the BM, we isolated the BM cells from non-diabetic (ND), 10-week diabetic (D), and diabetic mice treated with the specific LOX inhibitor β-aminopropionitrile (D+BAPN) and measured the protein expression levels of LOX and collagens IV and VI, key matrix proteins of the BM physical microenvironment. Western blot analysis revealed that diabetes leads to a significant increase (by 1.8 fold; p<0.05) in BM LOX expression, which was predictably associated with upregulation of BM collagen IV (by 2.5 fold; p<0.05) and collagen VI (by 1.7 fold; p<0.05) (**Fig. 1A**). Notably, this diabetes-induced upregulation of BM matrix proteins was blocked by LOX inhibitor BAPN. These findings were independently confirmed by immunostaining BM sections with anti-LOX, anti-collagen IV, and anti-collagen VI **(Fig. 1B)**.

**Figure 01:**
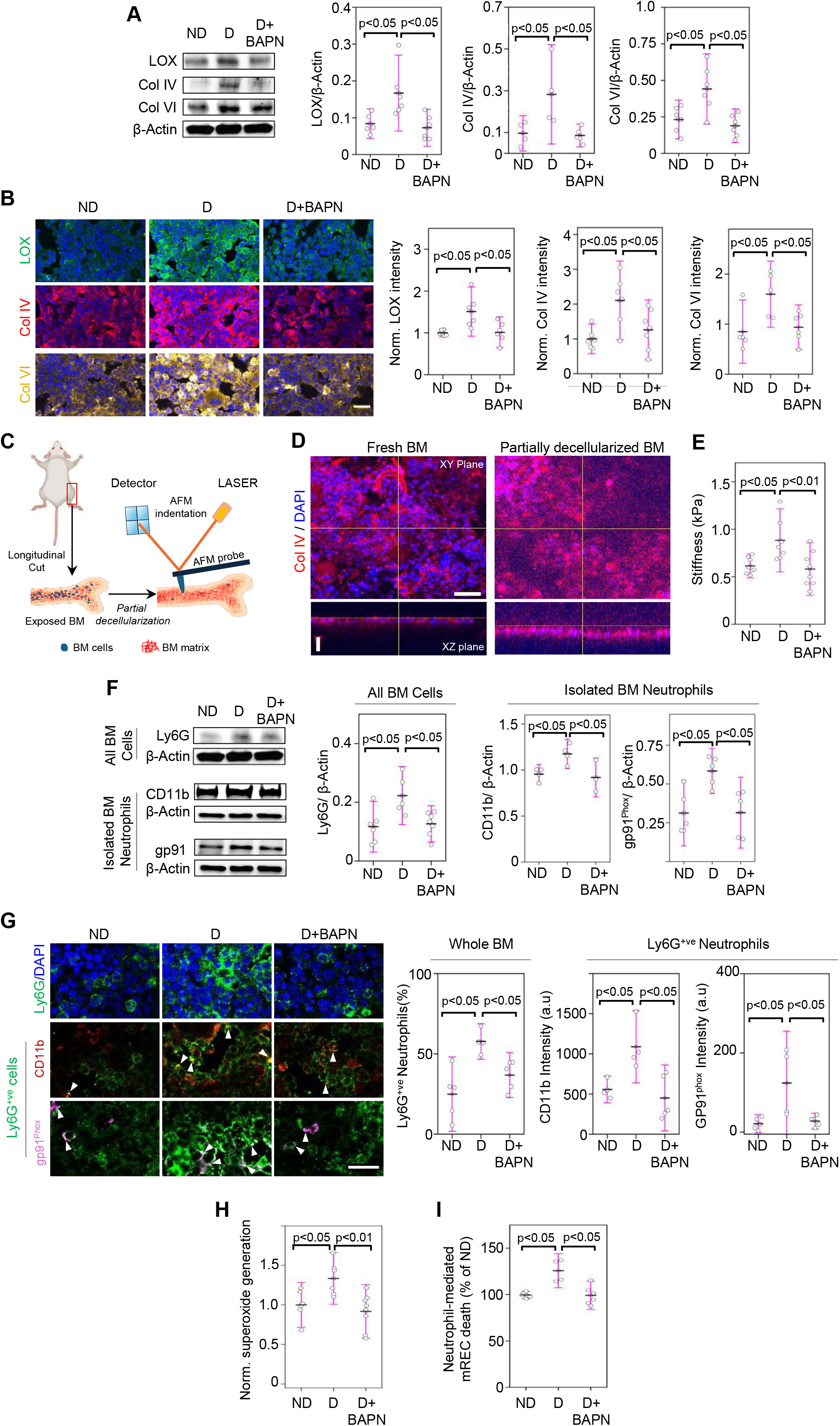
LOX promotes BM stiffening and neutrophil activation in diabetes. **(A)** Representative Western blot bands and cumulative densitometric analyses show normalized (w.r.t. β-actin) LOX, collagen IV, and collagen VI protein expression in BM cells isolated from nondiabetic (ND), 10-week diabetic (D), and diabetic mice treated with LOX inhibitor BAPN (D+BAPN; 3 mg/kg BW) (n=4-6 mice/group). **(B)** Representative immunofluorescence images and subsequent fluorescence intensity analysis reveal LOX (*green*), collagen IV (*red*), and collagen VI (*cyan*) levels in BM sections obtained from ND, D, and D+BAPN mice (n=5–6 mice/group). Scale bar: 20 µm. **(C)** Schematic shows BM preparation for AFM stiffness measurement. **(D)** Representative confocal images in XY and XZ planes show a typical mouse BM labelled with anti-col IV (red) and DAPI (blue) before and after partial decellularization. **(E)** Data plot indicates AFM stiffness measurement of the superficial BM matrix obtained from ND, D, and D+BAPN mice (n=7-8 mice/group). Data was obtained from ≥30 locations per BM. Scale bar: 50 µm. **(F)** Representative Western blot bands and cumulative densitometric analyses show normalized (w.r.t. β-actin) Ly6G protein levels in whole BM cells and CD11b and gp91^Phox^ levels in isolated BM neutrophils from ND, D, and D+BAPN mice (n=4-7 mice/group). **(G)** Representative immunofluorescence images of BM sections and subsequent fluorescence intensity analysis reveal the percentage Ly6G^+ve^ neutrophils (magenta) among all (DAPI-labeled) BM cells as well as the CD11b (green) and gp91^Phox^ (yellow) levels in the Ly6G^+ve^ neutrophil fraction. Scale bar: 20 µm. **(H)** Data plot shows superoxide generation by DHE-labeled neutrophils isolated from ND, D, and D+BAPN mice (n=6-7 mice/group) following brief fMLP stimulation. Data were normalized w.r.t. ND condition. **(I)** Flow cytometric analysis of FITC-Annexin V-labeled mRECs reveal endothelial apoptosis following 16h co-culture with neutrophils isolated from ND, D, and D+BAPN mice (n=4-6 mice/group). All data plots indicate mean±SD from multiple mice.

LOX-mediated increase in collagen deposition and crosslinking is known to stiffen retinal subendothelial matrix and tumor stroma. To determine whether LOX similarly increased BM matrix stiffness in diabetic mice, we harvested mouse femurs, cut them longitudinally to expose the BM, and subjected them to partial decellularization to remove the superficial layers of cell, thereby exposing the BM matrix **(Fig. 1C)**. Labeling the partially decellularized BM with anti-collagen IV and DAPI confirmed the substantial depletion of BM cells (as judged by marked reduction in DAPI staining) while retaining the collagen IV matrix **(Fig. 1D)**. Importantly, AFM stiffness measurement of the partially decellularized BM revealed that diabetes leads to a significant increase (by ∼1.5 fold; p<0.05) in BM matrix stiffness, which is blocked by LOX inhibition **(Fig. 1E)**. To our knowledge, this study is the first to demonstrate altered bone marrow stiffness in diabetes and implicates LOX as the key driver of this mechanical phenotype. Further, this finding raises the possibility that other systemic conditions marked by LOX upregulation, such sepsis and aging (9, 10), also exhibit BM stiffening.

Extracellular matrix stiffness is known to regulate cell behavior via mechanotransduction, the process by which mechanical cues (such as stiffness) are transduced into intracellular biochemical signaling (11). Thus, we asked whether diabetes-induced BM matrix stiffening alters the behavior of resident neutrophils, key determinants of retinal EC death associated with early diabetic retinopathy. To address this question, we first assessed BM neutrophil density by measuring the expression levels of neutrophil marker Ly6G in the whole BM cell population. Western blot analysis revealed a significant increase (by ∼two fold; p<0.05) in Ly6G expression in BM cells isolated from diabetic mice, thus indicating an increase in neutrophil numbers (**Fig. 1F**). This increase was blocked by LOX inhibitor BAPN. Importantly, the BM neutrophils were also significantly activated in diabetes, as judged by the higher protein expression levels of activation markers CD11b (a leukocyte integrin subunit that promotes EC binding) and gp91^phox^ (a subunit of NADPH oxidase complex that produces superoxide) in isolated BM neutrophils **(Fig.1F)**. Significantly, LOX inhibition blocked this diabetes-induced increase in BM neutrophil activation. This LOX-dependent increase in BM neutrophil density and activation in diabetic mice was independently confirmed by immunostaining BM sections with anti-Ly6G, anti-CD11b, and anti-gp91^phox^ **(Fig. 1G)**.

Neutrophil activation is implicated in retinal EC apoptosis associated with early diabetic retinopathy (2, 6). Consistent with this, activated BM neutrophils isolated from diabetic mice exhibited higher levels of (cytotoxic) superoxide release than their nondiabetic counterparts **(Fig. 1H)**, which agreed well with their higher expression of gp91^phox^ **(Fig. 1G)**. This superoxide overproduction by diabetic BM neutrophils was associated with an ∼25% increase (p<0.05) in the apoptosis of co-cultured mouse retinal ECs **(Fig. 1I)**. Crucially, LOX inhibition suppressed the diabetes-induced increase in neutrophil superoxide generation and cytotoxicity towards retinal ECs.

### LOX and BM matrix stiffening synergistically activate resident neutrophils

LOX exists in both matrix-bound and soluble forms. While the matrix-bound LOX regulates cell behavior via matrix stiffening and altered mechanotransduction, soluble LOX can exert direct biochemical effects on cells (7, 12). To determine the relative contribution of BM matrix-bound LOX, and hence BM matrix stiffness, to diabetes-induced neutrophil activation and cytotoxicity, we developed BM matrix-mimicking gelatin methacrylate (Gel-MA) hydrogels whose stiffness can be tuned by adjusting the concentration of photocrosslinker LAP. Rheological measurements indicated that LAP concentrations of 0.08% and 0.14% (w/v) yield hydrogel stiffness (normal’-0.5 kPa; ‘stiff’-1.8 kPa) comparable to that of ND and D mouse BM matrix, respectively **(Fig. 2A**).

**Figure 02:**
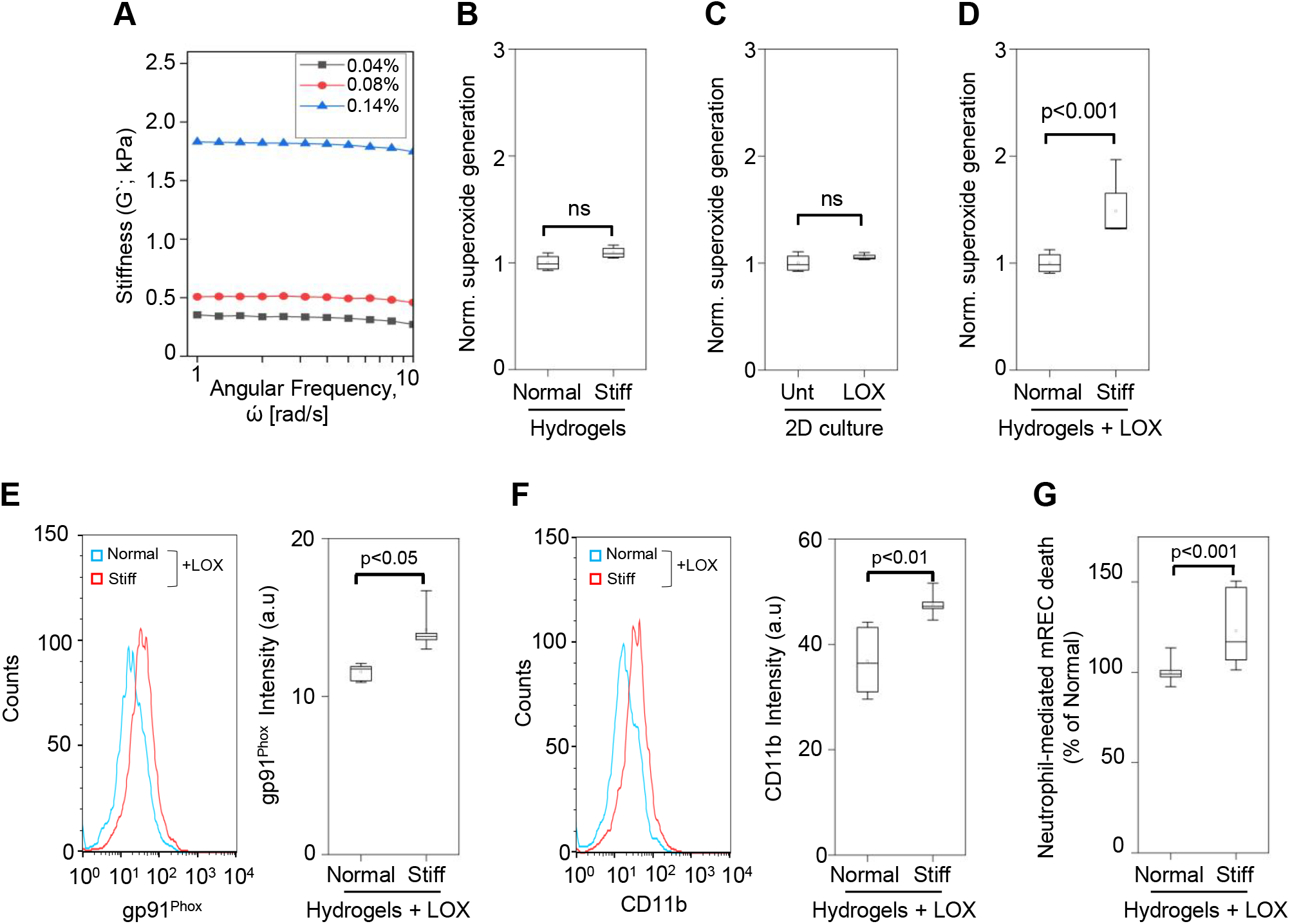
LOX and BM stiffening synergistically activate resident neutrophils. **(A)** Graph plot shows the bulk stiffness measurement (storage modulus, G’) of 5% (w/v) Gel-MA hydrogels prepared with varying photocrosslinker (LAP) concentration (0.04-0.14%) and subjected to a frequency sweep using an oscillatory shear rheometer. **(B)** Box whisker plots show superoxide generation by mouse neutrophils extracted from ‘normal’ or ‘stiff’ hydrogels following 6h culture. Data were normalized w.r.t. ‘normal’ condition. **(C)** Box whisker plot shows superoxide generation by mouse neutrophils grown in regular 2D cultures in the absence or presence of recombinant LOX (75 ng/mL; 6h). Data were normalized w.r.t. untreated condition. **(D)** Box whisker plots show superoxide generation by mouse neutrophils extracted from ‘normal’ or ‘stiff’ hydrogels following 6h culture in the presence of recombinant LOX (75 ng/mL). Data were normalized w.r.t. ‘normal’ condition. **(E, F)** Flow cytometric analysis (histogram) reveal differential gp91^phox^ (E) and CD11b (F) protein expression in mouse neutrophils extracted from ‘normal’ or ‘stiff’ hydrogels following 6h culture with recombinant LOX. (**G)** Flow cytometric analysis of FITC-Annexin V-labeled mRECs reveal endothelial apoptosis following 16h co-culture with neutrophils extracted from normal or stiff hydrogels. Box whisker plots show cumulative analysis of 5,000 neutrophils (in B-F) or 10,000 mRECs (in G). Black line and square within the box represent the median and mean, respectively.

Culturing mouse neutrophils within these 3D constructs revealed that hydrogel stiffening alone fails to enhance superoxide generation **(Fig. 2B)**. However, addition of soluble LOX, which independently did not activate neutrophils in 2D cultures **(Fig. 2C)**, led to a significant increase (by 1.5 fold; p<0.001) in superoxide production by neutrophils in stiff hydrogels (**Fig. 2D**). Predictably, this superoxide overproduction was associated with higher expression levels of neutrophil gp91^phox^ **(Fig. 2E)** and CD11b **(Fig. 2F)** as well as increased apoptosis of mouse retinal ECs co-cultured with neutrophils extracted from the stiff hydrogels (**Fig. 2G**). Together, these findings indicate that the matrix-localized and soluble forms of LOX act synergistically to mechano-chemically activate BM neutrophils and render them cytotoxic towards retinal ECs. Whether prolonged diabetes (beyond the 10 weeks’ duration studied here) leads to further BM stiffening such that the stiffer BM alone (independent of soluble factors such as LOX) can activate neutrophils remains to be determined.

Diabetes is known to induce biochemical, cellular, and anatomical defects in the BM niche that promotes dysfunction of resident stem/progenitor and stromal cells implicated in hematopoiesis and tissue regeneration (13, 14). The current study complements these findings by revealing that diabetes also induces LOX-dependent physical defects in the BM environment that leads to stiffness-dependent activation of resident neutrophils, a key determinant of retinal EC death and capillary degeneration in early diabetic retinopathy. It is possible that the diabetes-induced BM stiffening and neutrophil activation we observe contributes to the dysfunction of resident BM stem/progenitor/stromal cells. This idea is supported by a recent study where age-related BM stiffening was found to disrupt the hematopoietic stem cell niche that is required for stem cell maintenance and differentiation (15). Our findings also provide rationale to investigate the mechanotransduction pathways underlying the stiffness-dependent activation of BM neutrophils. Success in this pursuit may help uncover new classes of anti-inflammatory targets for diabetic complications.

## Material and Methods

A detailed description of the methods is provided as *Supporting Information*.

### Experimental Animals

All animal procedures were approved by University of California Los Angeles Institutional Animal Care and Use Committee. Diabetes was induced in adult male mice using streptozotocin and some diabetic mice were treated with LOX inhibitor β-aminopropionitrile (BAPN), as described in *Supporting Information*.

### BM Stiffness Measurements

Isolated femurs were cut open longitudinally and the BM was partially decellularized to expose the superficial BM matrix for stiffness measurement using NanoWizard^®^ 4 XP BioScience atomic force microscope (AFM; Bruker Nanotechnologies, Santa Barbara, CA) fitted with a pre-calibrated LC probe (Bruker AFM Probes, CA, USA), as described in *Supporting Information*.

### Statistics

Data analyses were performed using OriginPro2022 software (Origin Lab Corporation, USA) and statistical differences assessed using one-way ANOVA followed by Tukey’s or Dunnett’s *post hoc* multiple comparisons test. Studies comparing two experimental groups were subjected to two-tailed unpaired Student’s *t* test. Results were considered significant if *p*<0.05.

## Acknowledgments

This work was supported by NIH/NEI grant R01EY028242 (to K.G.), Jules Stein Eye Institute and Research to Prevent Blindness (RPB) Innovation Award (to K.G.), and RPB Unrestricted Grant to the Department of Ophthalmology at the University of California, Los Angeles. The authors thank Prof. Deming Sun for his insightful comments.

## Author Contributions

M.A. conceived the idea, designed and performed experiments, analyzed data, and wrote the manuscript, S.C. performed experiments and analyzed data. V.T. performed experiments, S.S. designed experiments and analyzed data, K.G. conceived the idea, designed experiments, analyzed data, and wrote the manuscript.

## Notes

### Competing Interest Statement

The authors have declared no competing interest.

