## Supplemental Information for "Lysyl oxidase-dependent bone marrow stiffening promotes neutrophil activation in diabetes"

Supporting Information

### Extended material and methods

#### Experimental Animals

All animal procedures were performed in accordance with the Association for Research in Vision and Ophthalmology (ARVO) Statement for the Use of Animals in Ophthalmic and Vision Research and approved by University of California Los Angeles Institutional Animal Care and Use Committee. Diabetes was induced in adult (8-week-old) male C57BL/6J mice (Jackson Laboratory, USA) by injecting freshly prepared streptozotocin solution (STZ; MP Biomedicals, USA; 60 mg/kg body weight in 10 mM citrate buffer; pH 4.5) intraperitoneally (i.p.) daily for five consecutive days. Mice were classified as diabetic if their fasting blood glucose was >275 mg/dL two weeks after the last STZ injection, with this time point being considered as the onset of overt diabetes. Age-matched C57BL/6J mice that only received citrate buffer served as nondiabetic controls. Some diabetic mice were treated with a specific and irreversible LOX inhibitor β-aminopropionitrile (BAPN; 3 mg/kg through drinking water; Catalog no. A3134-5G; Sigma-Aldrich, USA) for 10 weeks (from the onset of overt diabetes) prior to euthanasia and collection of bone marrow neutrophils, femurs, and tibia for further analyses. Nondiabetic, diabetic, and diabetic mice receiving BAPN are henceforth referred to as ND, D, and D+BAPN mice.

#### Isolation of Bone Marrow Cells and Neutrophils

Femur and tibia were harvested from euthanized mice and flushed with PBS to extract the BM. The isolated BM cell suspension was pelleted by centrifugation at 300 *g* for 10 min before resuspending the cell pellet in RBC lysyl buffer (catalog no. 20110, Stemcell Technologies, Canada) for 10 min. Finally, the RBC-free cell suspension was centrifuged at 300*g* for 10 min to obtain the pellets with total BM cells. To isolate neutrophils from the BM cell population, the EasySep™ Mouse Neutrophil Enrichment Kit (Catalog no. 19762, Stemcell Technologies, Canada) was used as per manufacturer's protocol, which results in >90% pure neutrophils.

**Western Blot**

To determine the protein expression levels of LOX, Collagen IV, Collagen VI, and Ly6G in the BM and CD11b and gp91$^{Phox}$ expression in BM neutrophils, equal amounts (20 or 50 μg) of total protein from the entire BM cell population or neutrophils were separated in 4-15% Mini-PROTEAN® TGX precast protein gels (Biorad, USA) and transferred to nitrocellulose membrane before probing with anti-LOX (Catalog no. NB100-2527, Novus Biologicals, USA), anti-Collagen IV (catalog no. NBP126549, Novus Biologicals, USA), anti-Collagen VI (Catalog no. OB136001, Southern biotech, USA), anti-Ly6G (Catalog no. NBP2-00441, Novus Biologicals, CA, USA), anti-CD11b (Catalog no. NB110-89474, Novus Biologicals, CA, USA), anti-gp91$^{phox}$ (catalog no. SC-1305435; Santa Cruz Biotechnology, USA), and anti-β Actin (loading control; Catalog no. A00702S; Genscript, USA), followed by appropriate secondary antibody conjugated to horseradish peroxidase (Catalog no. PI-1000 and PI-2000; Vector Laboratories). Protein bands were detected using a SuperSignal™ West Dura Extended Duration Substrate (Catalog no. 34075; Thermo Fisher Scientific, USA) and imaged using ChemiDoc XRS+ System (Biorad). Targets were assessed by technical duplicates obtained from three independent experiments. Densitometric analysis was performed using ImageJ.

**Bone Marrow Sectioning and Immunolabeling**

Bone sections were obtained as previously described (1). Briefly, the isolated mouse femurs were fixed with 4% (w/v) paraformaldehyde for 24 h at 4$^{o}$C prior to decalcification in 0.5 M EDTA (pH 7.4) for 48 h at 4$^{0}$ C under constant agitation. Next, the decalcified bone was kept in cryoprotectant solution composed of 20% (w/v) sucrose and 2% (w/v) polyvinylpyrrolidone (PVP) (in PBS) for 24 h at 4$^{o}$C before embedding in 8% (w/v) gelatin, 20% (w/v) sucrose, and 2% (w/v) polyvinylpyrrolidone (PVP) (in PBS) and storage at -80 $^{o}$C. 10 μm-thick BM sections were obtained from the embedded/frozen bones using a cryostat (Cryostat NX70, Thermo Fischer scientific, USA) and stored at -80$^{o}$C. For immunolabeling, the thawed BM sections were rinsed with PBS thrice for 5 min each, permeabilized and blocked with 1% (w/v) BSA and 0.1% (w/v) Triton-X in PBS for 30 min, and incubated with anti-LOX, anti-collagen IV, anti-collagen VI, anti-LY6G, anti-cd11b and anti-gp91$^{phox}$ in 1%BSA and 0.1% Triton-X (in PBS) for overnight at 4°C. Next, tissue sections were rinsed thrice 5 min each with PBS prior to incubation with an appropriate Alexa Fluor-conjugated secondary antibody (Jackson Labs) in 1% BSA (in PBS) for 1 h at RT. Immunolabeled cells were rinsed thrice with PBS, labeled with DAPI (to stain nucleus) and mounted with Fluoromount-G (Catalog no. 4958-0, Invitrogen, USA) for imaging using a confocal microscope (Zeiss LSM 770; 63X/NA 1.4 objective).

**Bone Marrow Stiffness Measurement**

Freshly-isolated mouse femurs were cut open longitudinally and the BM was partially decellularized by incubating the sliced bone in PBS for 2-3 h to expose the superficial BM matrix. BM stiffness was measured using the NanoWizard® 4 XP BioScience atomic force microscope (AFM; Bruker Nanotechnologies, USA) fitted with a pre-calibrated PFQNM-LC-A-CAL probe (spring constant ~0.1 N/m) containing a parabolic tip (70 nm radius). The AFM was coupled to a Zeiss Axiovert phase contrast microscope to facilitate sample visualization and measurement. Stiffness was measured in contact mode force spectroscopy mode by applying a 200 pN (set point) indentation force. Force curves from multiple locations on the BM matrix (n≥30/condition) were analyzed using JPK Data Processing Software.

**Superoxide Generation**

Superoxide generation by mouse BM neutrophils was measured using DHE dye (Catalog no. D11347, Invitrogen, USA), as previously reported (2). Briefly, freshly isolated neutrophils were suspended in $Ca^{2+}$ buffer (136 mM NaCl; 4.7 mM KCl; 1.2 mM $MgSO_4$; 1.1 mM $CaCl_2$; 1.2 mM $KH_2PO_4$; 5 mM $NaHCO_3$; 5.5 mM Glucose; 20 mM HEPES in dd$H_2O$) containing 10 μM DHE dye for 30 min prior to brief stimulation with fMLP (10 nM; 5 min). Next, equal number of cells were added to a 96-well plate (quadruplicates/condition) for fluorescence measurement (518/606 nm) using a spectrometer (Molecular device Spectromax iD5). Superoxide generation by neutrophils extracted from gelatin methacrylate hydrogels was assessed as described above but without fMLP stimulation.

Supporting Information

### Cell Culture and Treatment

Mouse retinal endothelial cells (mRECs) were purchased from Cell Biologics Inc. (Catalog no. C57-6065, Cell Biologics, USA) and cultured in vendor recommended medium (Catalog no. M1168; Cell Biologics, USA). mRECs were used until passage 10. Freshly isolated mouse BM neutrophils were cultured in RPMI 1640 medium (Gibco™, Thermo Fisher Scientific) supplemented with 10% fetal bovine serum (FBS; Catalog no. SH30396.03, Hyclone, Logan, UT) and 1% Penicillin-Streptomycin (Pen-strep; Gibco™, Thermo Fisher Scientific).

### Neutrophil Cytotoxicity towards Retinal Endothelial Cells

Neutrophil cytotoxicity towards mRECs was assayed as previously described (3). Briefly, freshly-isolated mouse neutrophils or neutrophils extracted from gelatin methacrylate hydrogels were added to mRECs at a ratio of 1:3 for 16 h (in triplicates). Next, the co-culture was detached using trypsin and labeled with anti-mouse CD144 (Catalog no. 138005 Biolegends, San Diego, USA) to identify mRECs, together with Annexin V (Catalog no. BD 550911 BD biosciences, USA) and Propidium Iodide (Catalog no. 537059 Sigma-Aldrich, USA) to assess mREC death by flow cytometry. A total of 10,000 events were counted for each sample. Results were analyzed by Flow Jo 10.7.2.

### Gelatin methacrylate hydrogel preparation and characterization

Gelatin methacrylate hydrogel (Catalog no. 5272, Advanced Biomatrix, USA) was reconstituted at 5% (w/v) in RPMI culture media (Catalog no. 11879020, ThermoFisher scientific, USA,) supplemented with 5 mM glucose. Next, the visible light-sensitive photo initiator lithium phenyl-2,4,6-trimethylbenzoylphosphinate (LAP) was added to the hydrogels at different concentrations (0.08, 0.12%, or 0.14% v/v) before exposure to visible light (405 nm) for 3 min to induce photocrosslinking (gelation). The stiffness (bulk storage modulus, G') of gelatin methacrylate hydrogels was measured by oscillatory shear rheometry (operating in frequency sweep mode from 1 to 10 rad/sec) using an Anton Paar MCR 302 rheometer with a cone and plate geometry (diameter: 10 mm, cone angle: 2°).

### Neutrophil Culture in Gelatin Methacrylate Hydrogels

Neutrophils were homogenously suspended in the gelatin methacrylate-LAP hydrogel solution at a density of $1 \times 10^6$ cells/mL prior to photocrosslinking. Neutrophil culture medium was added to the cell/hydrogel constructs in the presence or absence of recombinant LOX (75 ng/mL; Catalog no. LS-G 15002-50 LS bio, MA, USA) for 3h. Next, neutrophils were extracted from the hydrogels by enzymatic digestion with collagenase (1 mg/mL; catalog no. C1-BIOC, Sigma Aldrich, USA) dissolved in HBSS with 3 mM $CaCl_2$ (4). Extracted cells were used for subsequent assays.
